# Automated screening by 3D light-sheet microscopy with high spatial and temporal resolution reveals mitotic phenotypes

**DOI:** 10.1101/2020.01.20.912659

**Authors:** Björn Eismann, Teresa G Krieger, Jürgen Beneke, Ruben Bulkescher, Lukas Adam, Holger Erfle, Carl Herrmann, Roland Eils, Christian Conrad

## Abstract

3D cell cultures enable the *in vitro* study of dynamic biological processes such as the cell cycle, but their use in high-throughput screens remains impractical with conventional fluorescent microscopy. Here, we present a screening workflow for the automated evaluation of mitotic phenotypes in 3D cell cultures by light-sheet microscopy. After sample preparation by a liquid handling robot, three-dimensional cell spheroids are imaged for 24 hours *in toto* with a dual inverted selective plane illumination (diSPIM) microscope with a much improved signal-to-noise ratio, higher imaging speed, isotropic resolution and reduced light exposure compared to a spinning disc confocal microscope. A dedicated high-content image processing pipeline implements convolutional neural network based phenotype classification. We illustrate the potential of our approach by siRNA knock-down and epigenetic modification of 28 mitotic target genes for assessing their phenotypic role in mitosis. By rendering light-sheet microscopy operational for high-throughput screening applications, this workflow enables target gene characterization or drug candidate evaluation in tissue-like 3D cell culture models.

## Introduction

The cell cycle with its highly conserved and tightly regulated phases plays a key role in cancer development and progression. Cell cycle alterations are a hallmark of human tumors, and many cell cycle proteins have oncogenic properties (*1*). Pharmacological or genetic modulation of mitotic oncogene expression is therefore a highly promising treatment approach.

To study dynamic biological processes such as the cell cycle *in vitro*, 3D cell cultures provide a niche microenvironment that replicates the *in vivo* tissue more closely than traditional 2D methods (*2*). Cellular and subcellular morphologies can thus be tracked in a physiologically relevant context, allowing the characterization of therapeutic target gene function and the evaluation of molecular perturbations. However, live fluorescent imaging of 3D tissue-like cell cultures with conventional laser scanning microscopes is problematic due to insufficient acquisition speed, low resolution in the Z direction, excessive light scattering within the tissue and high phototoxicity (*3*).

To overcome these challenges, recent advances in selective plane illumination microscopy (SPIM) or light-sheet microscopy provide imaging capabilities with increased acquisition speed, excellent optical sectioning, and high signal to noise ratio (*4–6*). Phototoxicity is reduced by separating excitation and detection axes, and exciting fluorophores in a single thin layer with a scanning Gaussian beam. SPIM thus enables the evaluation of phenotypes at the subcellular level in whole-spheroid or whole-organoid 3D cultures, with sufficient temporal resolution to visualize fast processes such as mitosis (*7, 8*).

While these features in principle make SPIM microscopes ideally suited to high-throughput or high-content screens, their distinct geometry and the large volumes of data generated pose new challenges for sample preparation as well as data processing and analysis (*9, 10*). Automated phenotype evaluation usually requires the delineation of imaged structures (segmentation) and their clustering into functional groups (classification) (*11*). Classical machine learning methods such as random forests (RF) employ a user-defined set of features to categorise structured input data (*12, 13*). More recently, deep artificial neuronal networks such as convolutional neuronal networks (CNN) have emerged as a promising alternative (*14*). They can use unprocessed images as input and achieve image classification without the need for predefined features, often resulting in superior performance (*15–18*), but require large annotated training data sets which limits their usability (*19*).

Here, we developed a high-throughput screening workflow for the automated evaluation of mitotic phenotypes in 3D cultures imaged by light-sheet microscopy, from sample preparation to quantitative phenotype description. By using commercially available technology, this workflow is reproducible and easily adaptable to different cell culture models or molecular perturbations. A liquid-handling robot executes automated sample perturbation and mounting. Light-sheet imaging is performed with a dual inverted SPIM (diSPIM) microscope, a commercially available upright light-sheet system enabling high-throughput imaging of standard 3D cell cultures at isotropic resolution. A dedicated high-throughput image processing pipeline optimized for the diSPIM acquisition geometry combines convolutional neural network-based cell cycle phase detection with random forest-based classification to quantify phenotypic traits. Using this approach, we detect mitotic phenotypes in 3D cell culture models following modulation of gene expression by siRNA knock-down or epigenetic modification. Our fully automated workflow thus adapts light-sheet microscopy for applications in high-throughput screening in 3D cell culture models.

## Results

### Light-sheet imaging screen for high-content mitotic phenotype quantification

To evaluate the applicability of SPIM for high-throughput screening of mitotic phenotypes in 3D cell culture, we used an MCF10A breast epithelial cell line (*20*) stably expressing H2B-GFP to label DNA throughout the cell cycle. MCF10A cells provide an established and widely used model for benign breast tumors, with single MCF10A cells developing into multicellular 3D spheroids over the course of several days when seeded into laminin-rich hydrogel (Matrigel) (*21*). We selected 28 mitotic target genes of interest for a high-throughput screen based on reported mitotic roles and a strong correlation (Pearson correlation > 0.5) or anti-correlation (Pearson correlation < −0.5) of gene expression with altered methylation levels at one or multiple CpGs in the promoter or a distant regulatory genomic region, respectively (Methods and Supplementary Table 1). Target gene knock-down by siRNA transfection enabled us to analyze the effects of altered expression of these cancer-related genes in MCF10A cells. For the siRNA screen, two different siRNA were chosen per gene of interest, and MCF10A H2B-GFP cells were transfected by solid-phase reverse transfection (*22*). INCENP was used as a positive knock-down control due to its known severe effects on mitosis (*23*), while non-coding siRNA was used as negative control.

To achieve automated sample preparation, we developed a protocol for a liquid handling robot, which mixes the pretreated cells with Matrigel and mounts them in small Matrigel spots in a defined grid on the imaging plate (Figure 1a). This approach not only minimized the Matrigel volume to 0.2 μl per spot, reducing cost, but also ensured reliable positioning of samples with minimal pipetting variations or human errors.

**Figure 1:**
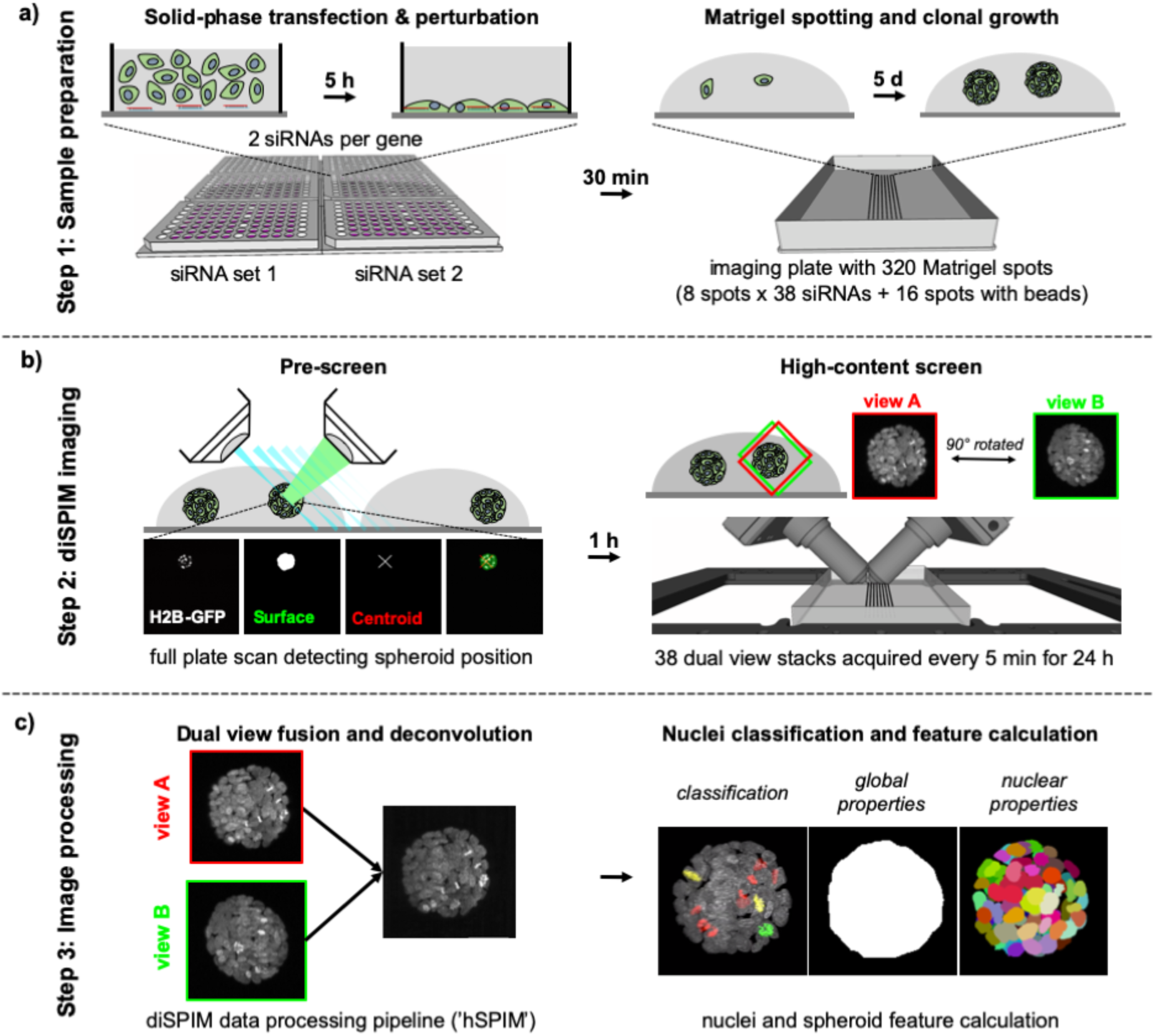
Light-sheet high-content live imaging screen. Key steps of the high-content screen. **a)** Step 1: Sample preparation. Cells were transfected in 2D by solid-phase reverse transfection in a 96-well plate format with two different siRNAs (siRNA set 1 and siRNA set 2) per target gene. Treated cells were then mixed with Matrigel and spotted in 0.2 μl droplets onto a one-well imaging plate. Over five days of culture, single cells clonally expanded into 3D spheroids. **b)** Step 2: diSPIM imaging. In a low-resolution stage scan pre-screen, positions of all spheroids were detected, and samples were selected for imaging by their position in the Matrigel spots; fused spheroids or cells growing on the plate in 2D were excluded. Subsequently, 38 individually treated samples were imaged every 5 minutes for 24 hours by dual view light-sheet microscopy, acquiring full stacks of view A (red) and view B (green) at a 90° angle to each other. **c)** Step 3: Data processing. Raw image data was processed by fusing visual information of view A and view B. Processed data was further analyzed to evaluate the phenotype of each spheroid throughout the time-lapse with regard to global spheroid features as well as single segment (single nucleus) properties.

After five days of cell culture at standard conditions, 3D spheroids were imaged with a diSPIM microscope for 24 hours at five-minute time intervals (Figure 1b). This choice of parameters accommodates the long cell cycle time of MCF10A cells (∼ 21 hours (*24*)), and enables single-cell tracking and the identification of subtle changes in the timing or morphology of nuclei undergoing mitosis.

Our imaging setup successfully realized the superior imaging capabilities of light-sheet microscopy over conventional confocal microscopes, including an improved signal-to-noise ratio, higher imaging speed, isotropic resolution and reduced light exposure compared to a spinning disc confocal microscope (Supplementary Table 2). Reductions in image quality due to in-depth light scattering were negligible after dual-view image fusion (Supplementary Figure 1). Due to the high temporal and spatial resolution, we were thus able to evaluate global, cellular and subcellular properties of mitotic gene knock-down phenotypes in live 3D spheroids (Supplementary Figure 2).

We acquired live long-term image data for three spheroids per siRNA, amounting to 74 terabytes of raw data for a total of 228 spheroids. To address the challenge of data processing (Figure 1c), we developed a high-throughput image processing tool called ‘hSPIM’, specifically tailored to the diSPIM geometry and acquisition properties. This pipeline allows for fast image fusion, deconvolution and data depth reduction of the raw diSPIM image data, based on a registration matrix and point spread function (PSF) detected from reference fluorescent beads. Sample background signal was reduced in our workflow by physically separating beads from spheroid samples; the registration matrix and PSF determined by imaging beads in Matrigel at a defined position was transferred to all acquired dual view stacks for registration and deconvolution. Additionally, we provide with ‘hSPIM’ basic single nuclei segmentation and textural feature detection. With most calculations executed in parallel on a high-performance GPU, we were able to process the raw data of one acquisition file within 8.23 seconds, reducing data size from 1.1 GB to 230 MB per position and time point (approximately 77 MB image data, 153 MB nuclei mask and 14 kB Haralick’s texture features) while significantly improving XYZ isotropic resolution and signal-to-noise ratio (Supplementary Figure 1).

The processed image data was subsequently analyzed in a KNIME (*25*) workflow. Key properties of individual spheroid development were quantified over time, including global descriptors of shape, volume and growth (e.g. total nuclei number, spheroid volume) as well as single nuclei specific traits such as cell cycle phase and position within the spheroid (Supplementary Table 3). To reliably identify the cell cycle phase of each nucleus, we compared the accuracy and performance of a random forest based approach with a VGG-based convolutional neuronal network classifier (*26*) applied to the same manually annotated nuclei set (Supplementary Figure 3). The random forest classifier relied on Haralick’s texture features (*27*) calculated by the ‘hSPIM’ processing pipeline and detected mitotic phase with 83% accuracy, whereas the CNN classified raw input image slices directly with 96% accuracy into the four key cell cycle stage (prophase, anaphase, metaphase, interphase) and was therefore chosen for all analyses.

### Detection of mitotic phenotypes in siRNA-treated MCF10A spheroids

Accurate cell cycle phase detection by CNN classification (Figure 2a) and the high temporal resolution of the diSPIM screen allowed us to track single cells through the different phases of the cell cycle (Figure 2b,c). In non-transfected control samples, 94.7% of all nuclei throughout the time lapse were classified as interphase, 0.9% as prophase, 1.2% as metaphase and 3.2% as anaphase (Figure 2d). As the high isotropic spatial resolution allowed us to determine nuclei and spheroid volumes at different stages, we also confirmed that prophase nuclei were on average the largest, followed by inter- and metaphase nuclei. While it has been suggested that nucleus position within the tissue influences cell cycle fate (*28*), we registered a similar radial distribution of cells in all cell cycle phases within MCF10A spheroids at day six of clonal development (Figure 2e).

**Figure 2:**
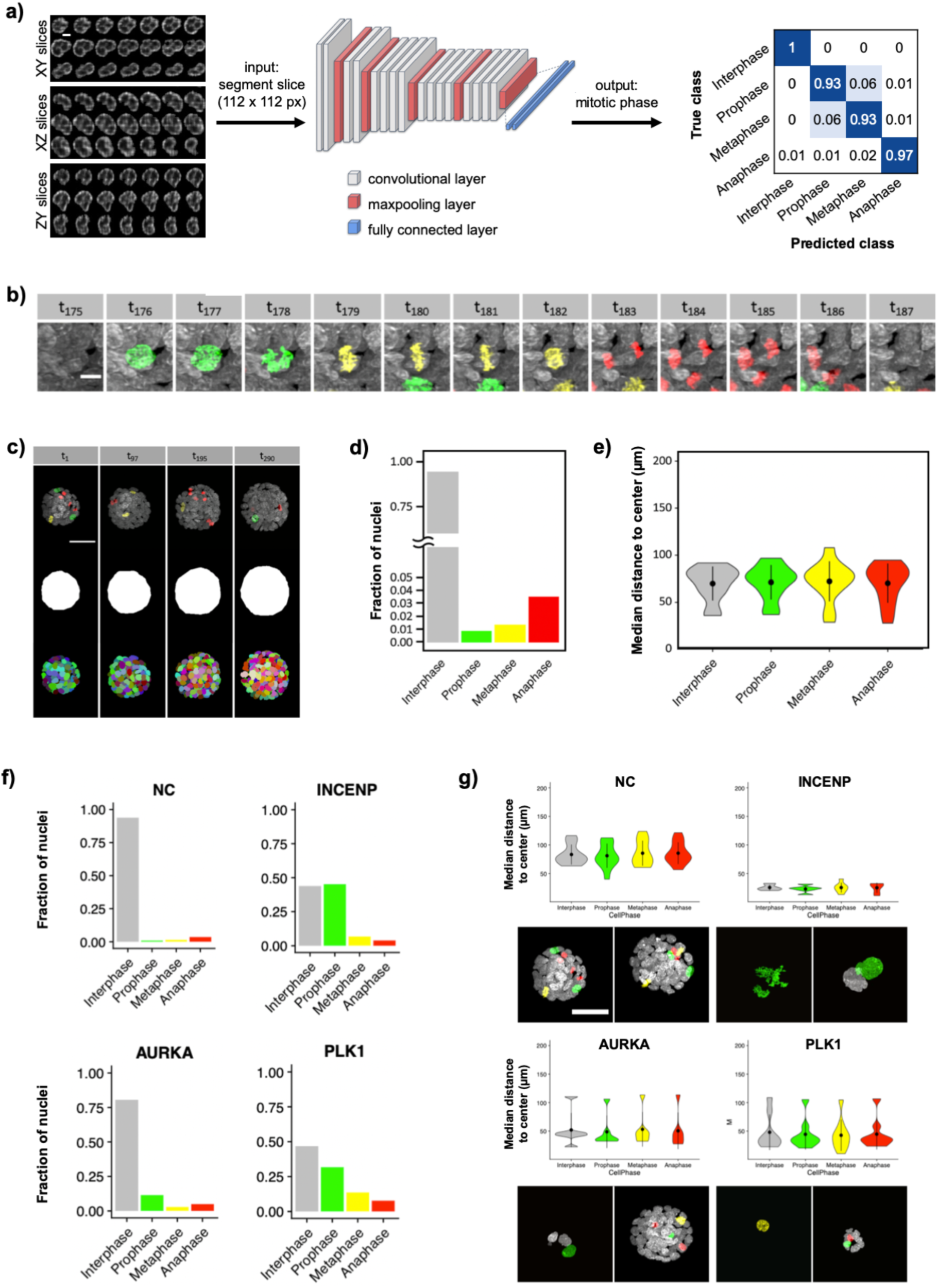
Image analysis of diSPIM data to detect mitotic phenotypes in 3D spheroids. **a)** For cell cycle phase detection, 2D image slices of the 3D segments were used as input to a VGG-based convolutional neuronal network consisting of convolutional, maxpooling and fully connected layers as indicated (see also Supplementary Figure 3). The network outputs a probability for each of the cell cycle phases, with cross correlation values shown as a measure of classification accuracy (cross correlation values represent 10% of the manually annotated training data set, with the other 90% used for training the network). **b)** Example time lapse (time points 175-187) of an untreated MCF10A cell undergoing mitosis, with interphase (white), prophase (green), metaphase (yellow) and anaphase (red) detected by deep learning image classification (scale bar = 5 μm). **c)** Four exemplary time points (t) of spheroid development imaged over 24 hours (time points 1-290), with the classified cells (colors as in **b**), spheroid hull and segment maximum projection displayed (scale bar = 50 μm). **d)** Bar plot showing the total fraction of control nuclei detected in different cell cycle phases throughout the screen (n = 205,068). **e)** Violin plot depicting the distance of nuclei from the spheroid center during the different cell cycle phases (n = 205,068). Black dots represent the median and whiskers the 25-75% interquantile range. **f)** Examples of abnormal mitotic phase durations induced by different siRNAs. Bar plots show the total fraction of nuclei detected in different cell cycle phases for negative control samples transfected with non-coding siRNA (NC), as well as spheroids transfected with siRNAs against INCENP, AURKA and PLK1. **g)** Examples of spheroid growth defects depicted by abnormal nuclei positions. Violin plots show the median distance of cells from spheroid centers during the different cell cycle phases for the same spheroids as in **f**. Black dots represent the median and whiskers the 25-75% interquantile range. Images show representative maximum and minimum sized spheroids at the start of the time lapse acquisition (scale bar = 50 μm).

In siRNA transfected samples, we observed a wide range of mitotic phenotype alterations. An over-representation of different cell cycle phases compared to control MCF10A spheroids indicated cell cycle arrest; INCENP and AURKA knock-down spheroids, among others, showed more cells in prophase (Figure 2f), while MYC and ATOH8 knock-down resulted in more anaphase cells. PLK1 knock-down nuclei displayed an increase in all mitotic classes, suggesting an elongated cell cycle (Figure 2f). Apoptotic cells frequently led to the assignment of improper mitotic phase transitions (such as prophase to interphase) in PLK1, EME1 and CEP85 knock-down spheroids. Defects in spheroid growth were identified in INCENP, AURKA and PLK1 knock-down spheroids, among others, whereas we did not observe abnormal positioning of cells in individual cell cycle phases (Figure 2g).

Hierarchical clustering of all mitotic phenotype quantifications (Supplementary Table 3) distinguished five major phenotypic groups (Figure 3). Cluster 1 comprised spheroids with a high growth rate closely resembling the non-coding siRNA control. Cluster 2 mostly contained samples with increased nuclear volumes during prophase and a higher proportion of cells in this phase, indicating prophase arrest and formation of macronuclei with increased DNA content. A higher volume growth rate was detected in spheroids in cluster 3, with some knock-down target genes (LMNB2, F11R, LHFP) also resulting in larger nuclei during anaphase. Cluster 4 spheroids showed strong phenotypes with reduced spheroid volume growth and a low total number of cell cycle transitions, indicating diminished mitotic activity. Finally, cluster 5 spheroids showed aberrant phenotypes in several features, describing grave cellular and mitotic defects.

**Figure 3:**
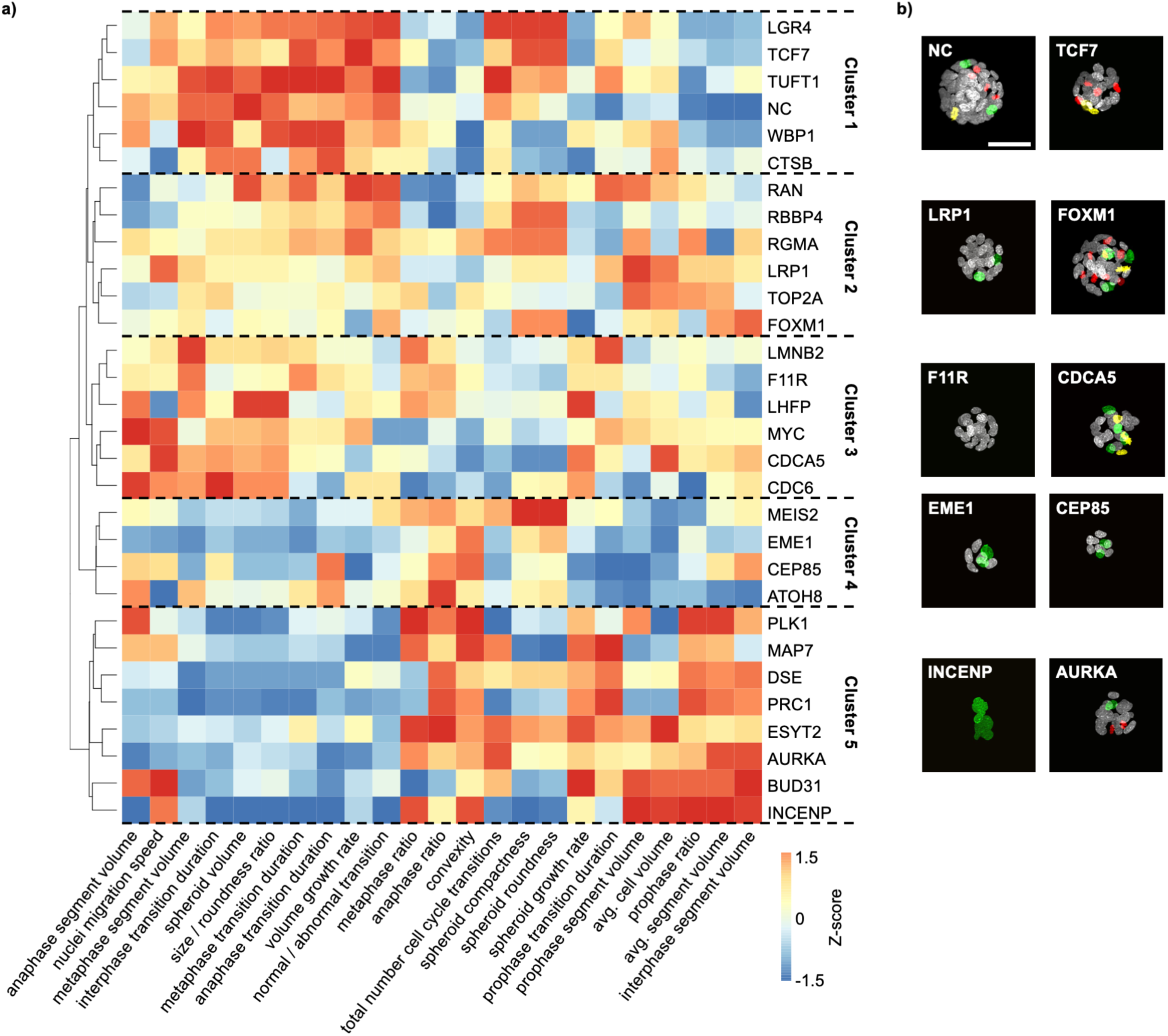
Clustered phenotype analysis of all features detected in diSPIM high-content screen. **a)** Rank-based hierarchical clustering of siRNA knock-down mitotic phenotypes by features describing the global and nuclei specific properties displays distinct clusters of siRNA target genes. **b)** Example images of spheroids from each cluster with classified nuclear mitotic phases (white, interphase; green, prophase; yellow, metaphase; red, anaphase). Scale bar = 50 *μ*m.

To evaluate the performance of the diSPIM screening workflow and analysis pipeline, we compared detected phenotypes to the MitoCheck database assembled from imaging the first two to four cell divisions of HeLa H2B-GFP cells upon siRNA target gene knock-down (*23*). Notably, despite the use of different cell lines, all of the strong phenotypes identified in our study were also described in the MitoCheck screen, including reduced spheroid growth in EME1 and ATOH8 knock-down cultures, elongated cell cycle phases in ESYT2 and PLK1 knock-down cells, as well as cell cycle arrest and increased apoptosis in INCENP, MAP7, DSE and PRC1 knock-down cells. Minor phenotypes such as macronuclei formation in some spheroids with CEP85 or MEIS2 knock-down and interphase arrest induced by MYC siRNA transfection were not detected by the MitoCheck screen.

### Detection of mitotic phenotypes after epigenetic perturbation

To further demonstrate the utility of our workflow, we used a novel molecular tool based on the CRISPR/Cas system (*29, 30*) to target regulatory epigenetic elements of the selected mitotic genes. Fusion proteins of deactivated Cas9 with no endonuclease activity (dCas9) with the effector domain of methylome-modifying proteins (dCas9-ED) have been shown to alter the epigenome at a targeted genomic location defined by an appropriate sgRNA (*31, 32*). We designed fusion proteins of dCas9 and the effector domain of either DNMT3a methyltransferase to achieve CpG methylation or TET1 for demethylation (Supplementary Figure 4). A dCas9 without added effector domain was used as a control physically blocking binding sites for regulatory factors. As MCF10A cells are highly resistant to plasmid transfection, we moved to human embryonic kidney 293 (HEK293) cells, which also develop into multicellular spheroids when seeded in Matrigel, and generated cell lines stably expressing the different constructs.

In a 2D pre-screen, we detected overall low effectivity of the dCas9-ED fusion proteins across our set of target genes (Methods and Supplementary Figure 5), but identified the target gene RGMA as robustly showing a mitotic phenotype under different methylome modifying conditions (dCas9-DNMT3a with sgRNAs targeting anti-correlated CpGs or the transcription start site (TSS), and dCas9-TET1 with sgRNAs targeting correlated CpGs). We therefore selected RGMA for 3D screening of mitotic phenotypes upon epigenetic modification, using the same sample preparation and imaging workflow as described above for the siRNA screen. Three and five days after sgRNA plasmid transfection, every sgRNA transfected spheroid (as assessed by GFP expression) was imaged and evaluated by Hoechst staining for mitotic phenotypes.

Most prominently, and in agreement with the MitoCheck database, macronuclei were detected in 41-52% of all spheroids transfected with either dCas9-DNMT3a and sgRNA targeting anti-correlated CpGs (Figure 4a), dCas9-TET1 and sgRNA targeting correlated CpGs (Figure 4b), or dCas9 and dCas9-DNMT3a with sgRNA locating to the TSS (Figure 4c,f). Transfection with dCas9 control protein and sgRNA targeted to regulatory CpGs resulted in macronuclei in only 7% (anti-correlated CpGs) or 6% (anti-correlated CpGs) of spheroids (Figure 4d,e), confirming the specificity of our epigenetic modification of RGMA. Reduced spheroid growth, apoptotic condensed DNA and elongated mitosis or mitotic arrest were also frequently observed, indicating that RGMA knock-down has severe effects on cellular homeostasis (Figure 4g).

**Figure 4:**
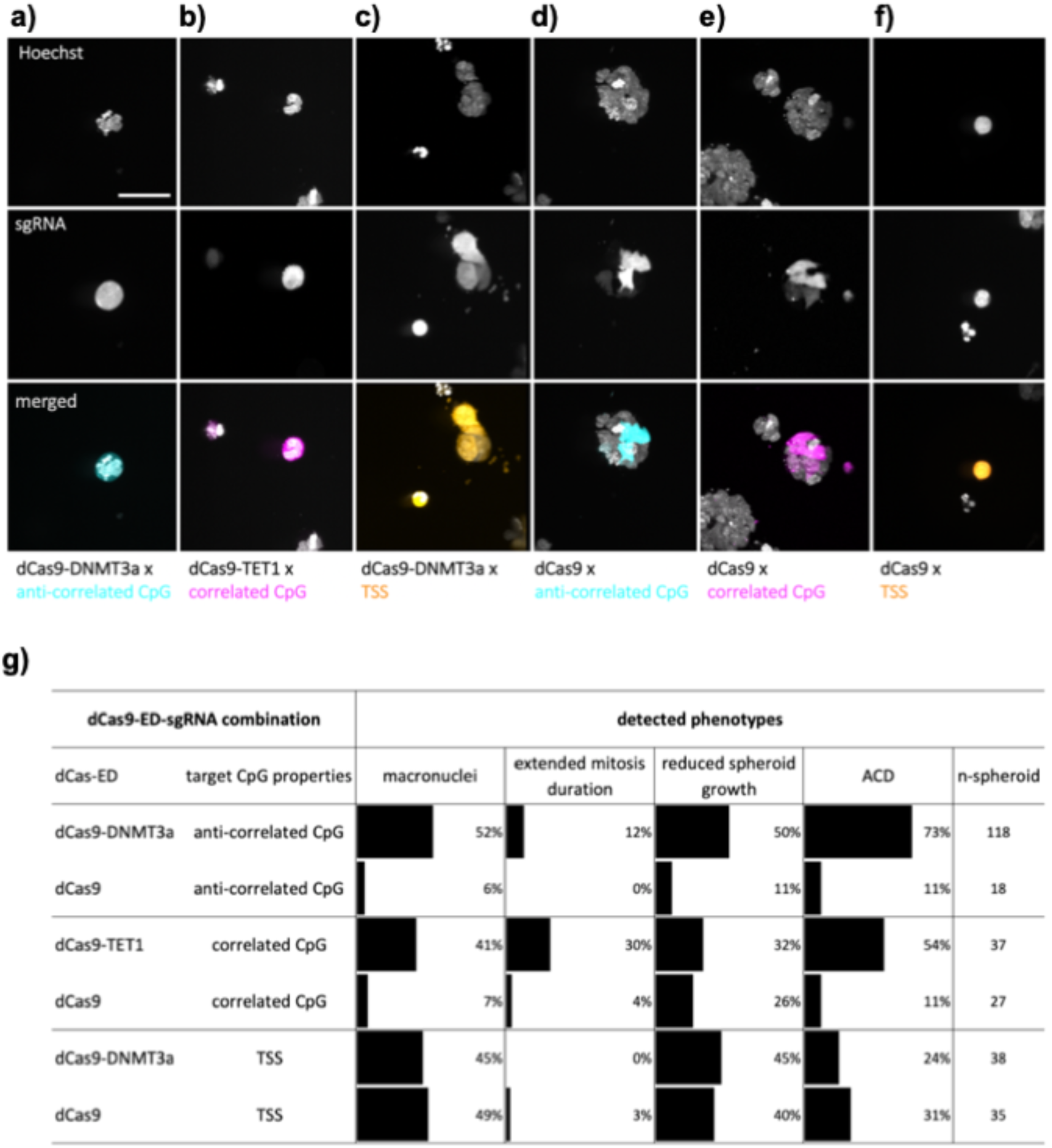
Phenotype analysis of dCas9-ED targeting RGMA regulatory CpGs in 3D HEK293 spheroids. Example images of HEK293 spheroids stably expressing dCas9-ED or dCas9, after transfection with sgRNA designed to epigenetically alter RGMA expression, in the following combinations: **a)** dCas9-DNMT3a with sgRNA targeting anti-correlated CpG (cyan), **b)** dCas9-TET1 with sgRNA targeting correlated CpG (magenta), **c)** dCas9-DNMT3a with sgRNA targeting the TSS (orange), **d)** dCas9 with sgRNA targeting anti-correlated CpG (cyan), **e)** dCas9 with sgRNA targeting correlated CpG (magenta), **f)** dCas9 with sgRNA targeting the TSS (orange). Scale bar = 50*μ*m. **g)** Summary of phenotypic effects of epigenetic targeting of RGMA expression in HEK293 spheroids. The percentage of spheroids displaying abnormal cellular and global properties, including macronuclei formation, extended mitosis duration, reduced spheroid growth and apoptotic condensed DNA (ACD), is shown.

Modulation of RGMA gene expression at the transcriptional level, using a dCas9-ED system, and at the translational level, using siRNA, thus results in similar mitotic phenotypes which can be detected with our high-throughput light-sheet imaging workflow, illustrating its versatility.

## Discussion

The high-throughput light-sheet live imaging workflow presented here provides a novel tool for screening individually treated 3D cell cultures with high spatial and temporal resolution, signal-to-noise ratio, fast acquisition speed and minimal phototoxic effects. Automation of the different steps from sample treatment and mounting to spheroid position detection and image acquisition, as well as the commercial availability of all materials, ensure that the workflow is reproducible and applicable to different culture models or treatment methods. Furthermore, we provide an easy-to-use image processing pipeline adapted to the geometry of the dual-view inverted light-sheet system, using a CNN for reliable cell cycle phase classification in 3D.

By applying this workflow to 3D cultured MCF10A H2B-GFP cells transfected with siRNA targeting mitotic genes, and HEK293 cells transfected with dCas9-based epigenetic modifiers, we were able to evaluate the effect of single gene knock-down on key cellular and spheroid features. Our integrated high-content analyses also highlight similar phenotypes caused by different genes. Due to the superior temporal and spatial resolution provided by the diSPIM system in combination with long-term acquisition, we could track cells over 24 hours and detect subtle mitotic 3D phenotypes not accessible with conventional fluorescent microscopy. As the diSPIM geometry uses dipping lenses, phenotypes that can be evaluated by high-throughput live imaging with this workflow are restricted to intracellular perturbations such as siRNA or CRISPR-Cas9 based screens, although an end-point analysis of fixed samples can also be conducted. To extend this workflow to other applications, such as using small molecule libraries for high-throughput drug screens, samples need to be physically separated into distinct wells during culture. Novel cell culture plate formats could accommodate this, for example by mounting spheroids on invertible pillar structures for culture in multiwell plates.

The light-sheet imaging setup presented here is thus adaptable for high-throughput screening of 3D cell cultures in a variety of settings. As spheroids can be recovered after imaging, it is also compatible with combined approaches correlating image data with other modalities, including single-cell genomic or transcriptomic analyses. We therefore expect that the ability to quantitatively evaluate 3D phenotypes in live cell cultures at high throughput will advance functional characterizations of dynamic cellular processes in tissue-like microenvironments, in cancer research and beyond.

## Author contributions

BE and CC conceived the study; BE, TGK, HE, RE and CC designed experiments; BE, JB and RB prepared, tested and conducted siRNA coating in 96-well plate; BE and JB developed Hamilton liquid handling Matrigel/cell spotting protocol; CH defined targets for methylome modification; BE conducted diSPIM imaging screen; BE and CC designed and wrote in collaboration with mbits imaging GmbH the ‘hSPIM’ software; BE designed the KNIME analysis workflow; BE and LA developed the deep learning classification model; BE and TGK analyzed phenotypes; BE, TGK and CC wrote the manuscript. All authors revised and approved the manuscript.

## Acknowledgements

We thank Jon Daniels (Applied Scientific Instruments) for extensive technical support on the diSPIM systems, Christian Dietz for support with the KNIME analysis pipeline, Katharina Jechow (Theoretical Bioinformatics, DKFZ) for technical laboratory support, and Markus Fangerau and Ingmar Gergel (mBits) for development and support of the ‘hSPIM’ library. This study was supported by the Helmholtz International Graduate School for Cancer Research and the iMed Program (Helmholtz Association). TGK was supported by a DKTK Postdoctoral Fellowship. The Advanced Biological Screening Facility was supported by the CellNetworks-Cluster of Excellence, Heidelberg University (EXC81).

