## Supplementary Materials for "Automated screening by 3D light-sheet microscopy with high spatial and temporal resolution reveals mitotic phenotypes"

1. Supplementary Figures 1-5
2. Supplementary Tables 1-3

#### 1. Supplementary Figures

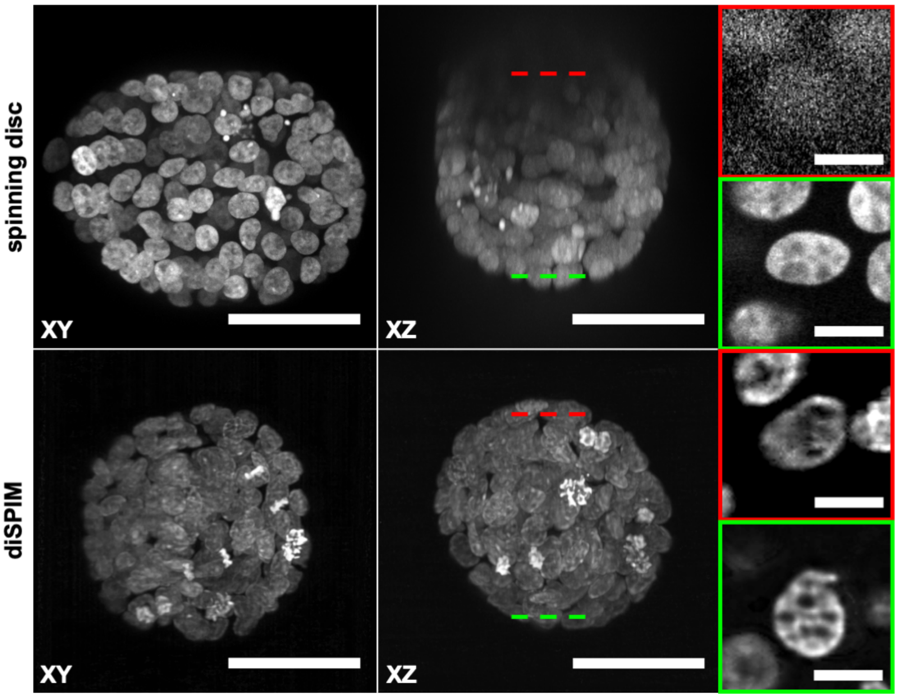


**Supplementary Figure 1: Comparison of spinning disc and light-sheet imaging of 3D spheroids**

Direct comparison of spinning disc and light-sheet imaging performance. MCF10A H2B-GFP spheroids with a size of about 80 µm in diameter were imaged six days after seeding single cells in Matrigel. XY and XZ maximum projections of the full 3D stack (scale bar = 50 µm) illustrate the XYZ resolution of the spinning disc and diSPIM microscopes. Inserts show single nuclei close to the detection objective (green box) or imaged 80 µm inside the sample (red box) in XY (scale bar = 10 µm), with the position of the corresponding Z-stack slices in the whole spheroid depicted by the red and green lines.

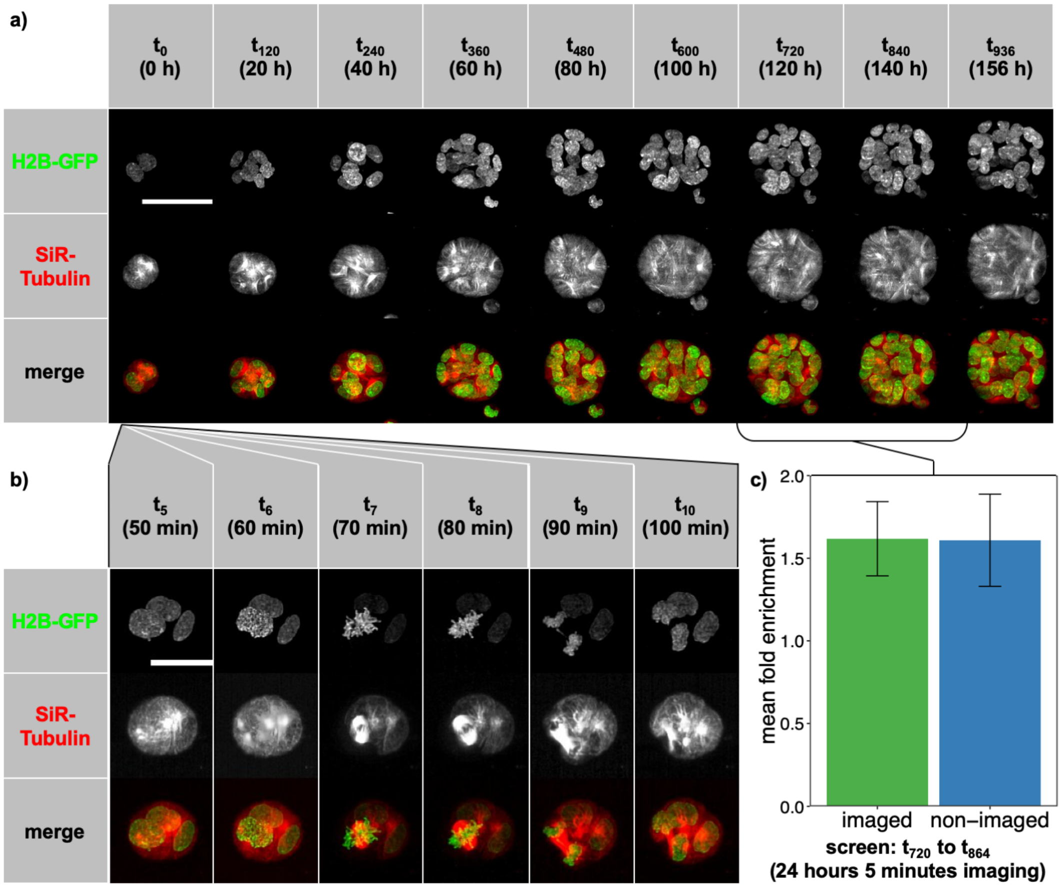


**Supplementary Figure 2: Long term imaging capabilities of the diSPIM**

**a)** Example of an untreated MCF10A H2B-GFP spheroid imaged over 156 hours / 936 time points (t) every 10 minutes from the two-cell stadium to the fully developed spheroid in two channels (H2B-GFP and SiR-Tubulin dye). Scale bar = 50 µm. **b)** High temporal and spatial resolution enable the detection of distinct features of the cytoskeleton and the different cell cycle stages (scale bar = 25 µm). **c)** Mean fold enrichment of the number of nuclei during a 24 hour acquisition cycle for imaged and non-imaged spheroids ($n_{imaged}$ = 31 / $n_{non\text{-}imaged}$ = 33). Error bars represent standard deviation.

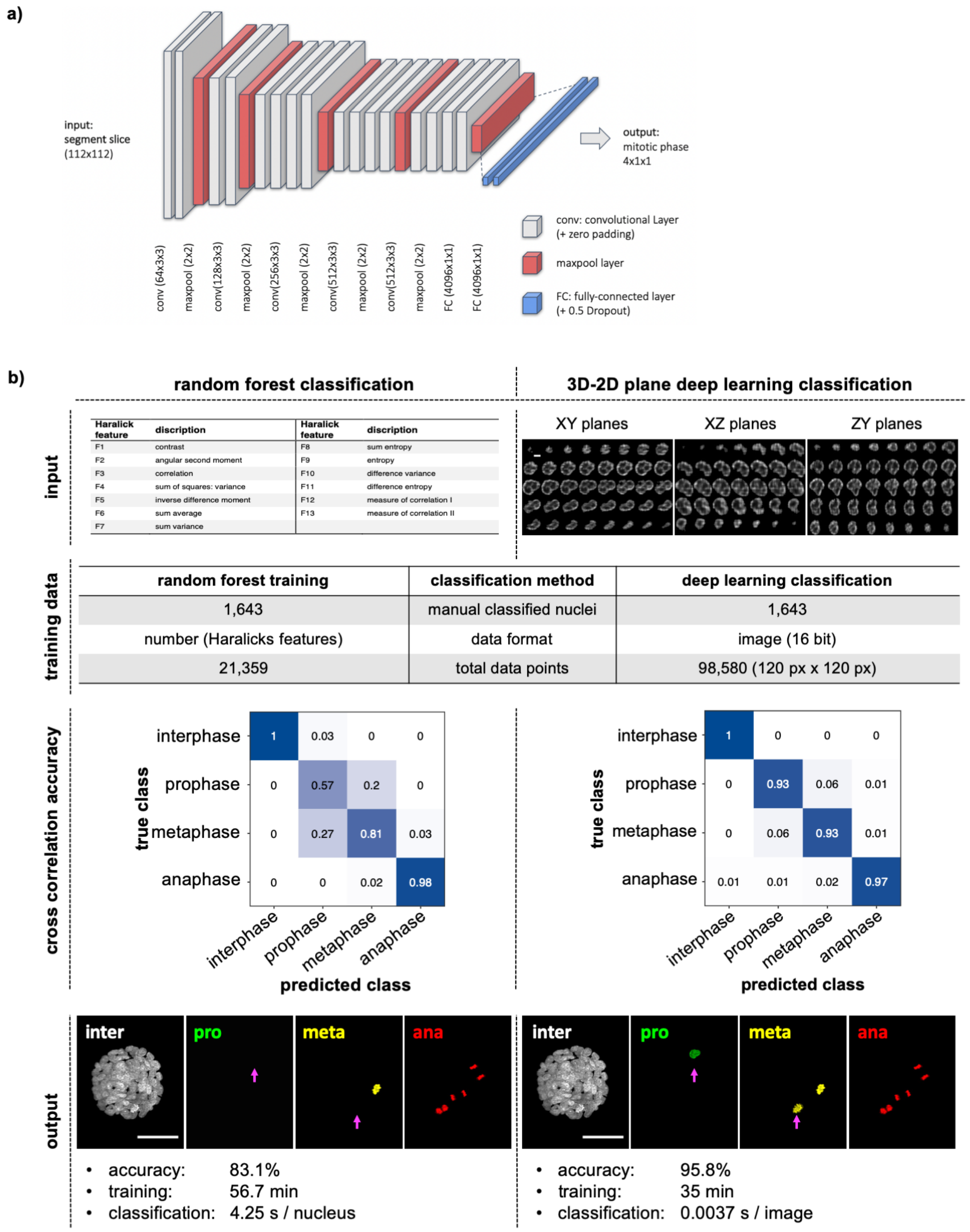


**Supplementary Figure 3: Comparison of mitotic cell phase detection by a Random Forest classifier versus a Convolutional Neuronal Network**

**a)** The VGG-based convolutional neuronal network uses 2D image slices of the 3D segments (112 px x 112 px) as input. The network consists of convolutional and maxpooling layers as indicated, with the output combined by two fully connected layers, and outputs a probability for each of the cell cycle phases. **b)** Comparison of the CNN with a Random Forest classifier. From top to bottom: Inputs were the 13 Haralick's features (F1-13) calculated by the 'hSPIM' data processing pipeline for random forest classification, and 2D (XY, XZ and YZ) slices of a 3D nucleus image for deep learning classification (scale bar = 5 µm). Manual labelling of the same nuclei resulted in a training data set comprising 21,359 data points for the RF classifier and 98,580 16-bit images for the CNN. Cross correlations are shown as measurements of classification accuracy. In direct comparison, classification differences between RF and CNN classification applied to the same image can be visually identified (magenta arrows; scale bar = 50 µm).

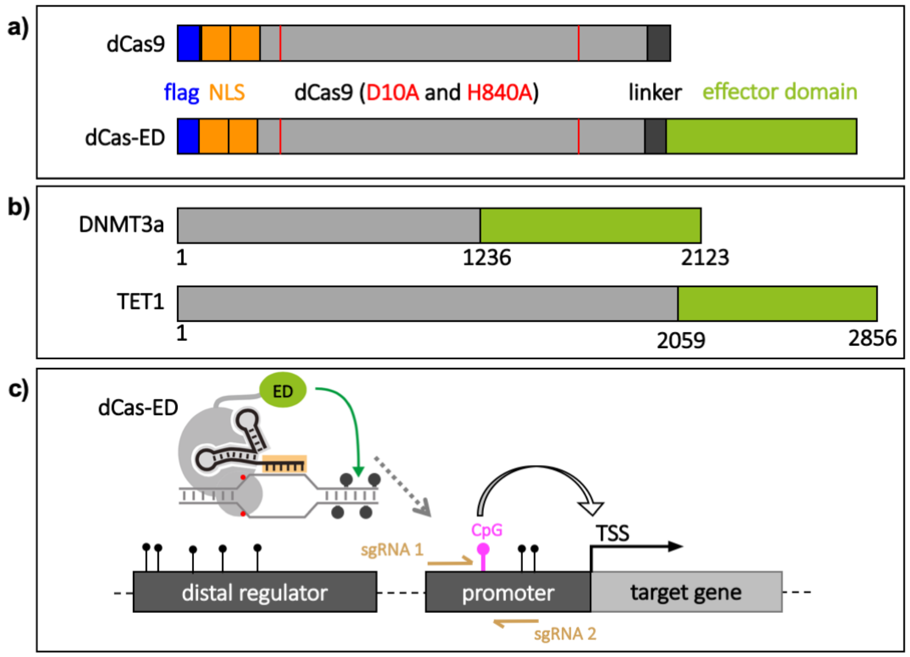


**Supplementary Figure 4: dCas9-ED targeting regulatory CpGs**

**a)** CRISPR-Cas9 based epigenetic modifiers and binding control were composed of an M2-flag (blue), two nuclear localization sequences (orange), the dCas9 mutated at residue 10 and 840 (red) deactivating the endonuclease function of the Cas9, and the effector protein (green). The effector domains were fused C-terminally via a linker (dark grey). **b)** Effector domains are the catalytically active, C-terminal domains of DNMT3A and TET1 from residue 1236 (DNMT3a) and 2059 (TET1) to the C-terminus of the protein. **c)** CRISPR-dCas9 fused with the effector domain was located to specific target sites defined by the sgRNA. Combinations of dCas9-ED with sgRNA targeting correlated or anti-correlated CpGs defined gene regulatory properties. Per CpG, two sgRNAs with opposite orientation were transfected, targeting loci upstream and downstream of the CpG (magenta). CpGs were located in promotors or distal regulatory regions.

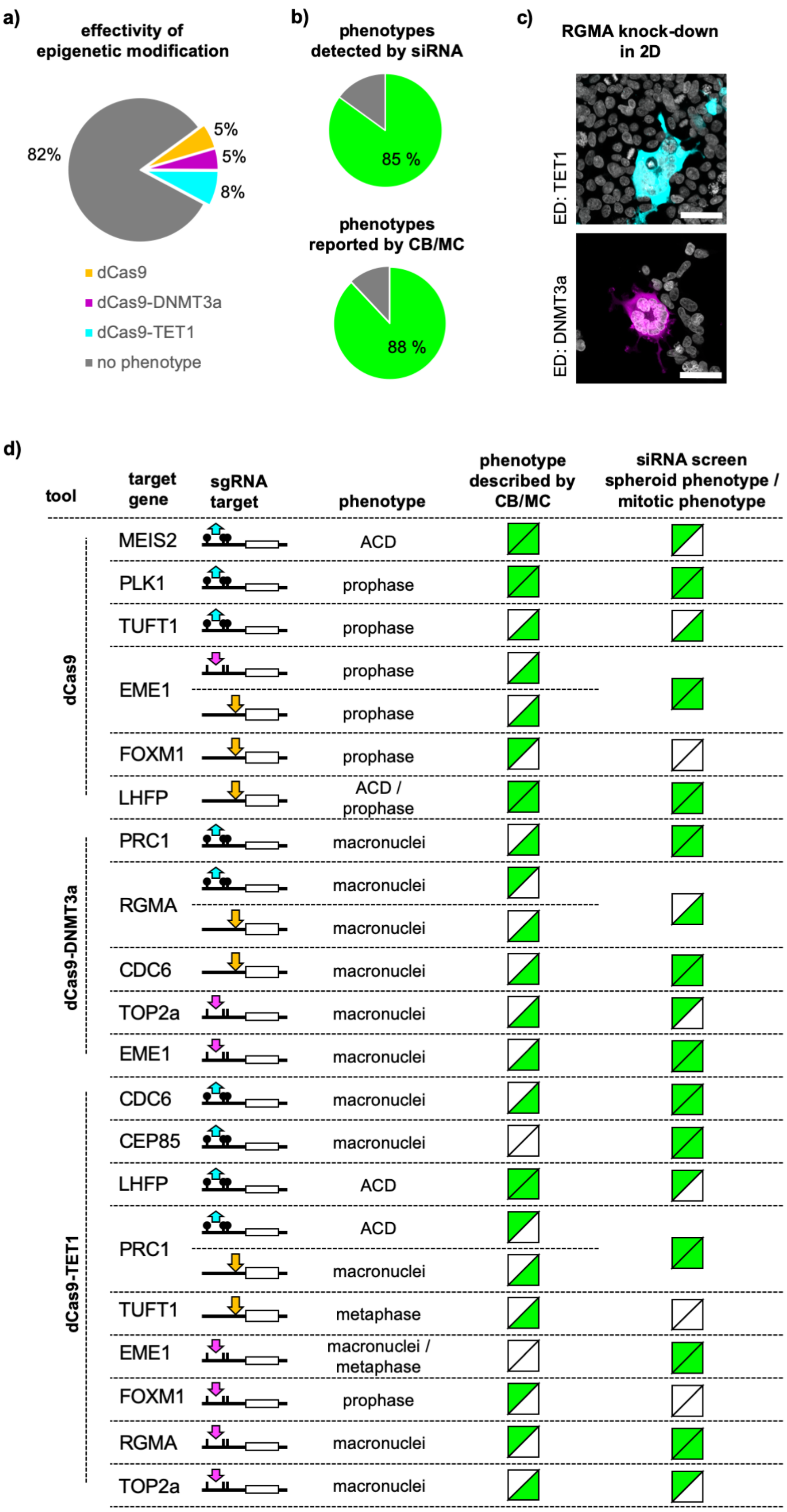


**Supplementary Figure 5: Comparative analysis of knock-down mitotic phenotypes**

**a)** Fraction of dCas9, dCas9-DMT3a and dCas9-TET1 combinations with sgRNAs targeting expression of the 18 genes shown in **d** that resulted in a mitotic phenotype in HEK293 cells cultured in

2D. **b)** Fraction of those phenotypes that was also detected in the siRNA screen (top) or reported in the Cyclebase (CB) and/or MitoCheck (MC) databases (bottom). **c)** Example images of mitotic phenotypes evoked in 2D HEK293 cells upon dCas9-ED localization to anti-correlated (cyan) or correlated (magenta) regulatory CpGs. **d)** List of target genes that showed a more than 1.5-fold increase in detection frequency of phenotypes (third column), such as increased representation of individual cell cycle phases or apoptotic condensed DNA (ACD), upon expression of dCas9 (top), dCas9-DNMT3a causing methylation (middle), or dCas9-TET1 causing demethylation (bottom) and transfection with sgRNA targeting anti-correlated (cyan) or correlated (magenta) regulatory CpGs or the transcription start site (orange) of the gene (second column). Detected phenotypes were compared with previously published databases (CB: Cyclebase, MC: MitoCheck) and siRNA diSPIM screen results. Green triangles indicate spheroid and mitotic phenotypes that were consistent across screens and sources.

#### 2. Supplementary Tables

##### Supplementary Table 1: Selected target genes

Target genes were selected based on their association with the cell cycle, and correlation of their expression with the methylation level of either correlated or anti-correlated CpGs with high average absolute Pearson correlation value ($R_{avg}$).

| Target gene | Name | Ambion siRNA # | Regulatory CpG | Relation of CpG methylation to gene expression | $R_{avg}$ |
|---|---|---|---|---|---|
| **ATOH8** | Protein atonal homolog 8 | s39645 / s39643 | 1 | anti-correlated | 0.52 |
| **AURKA** | Aurora kinase A | s196 / s197 | 2 | correlated / anti-correlated | 0.52 |
| **BUD31** | Protein BUD31 homolog | s17010 / s17009 | 1 | correlated | 0.45 |
| **CDC6** | Cell division control protein 6 | s2744 / s2746 | 2 | anti-correlated | 0.65 |
| **CDCA5** | Sororin | s41424 / s41425 | 6 | correlated | 0.70 |
| **CEP85** | Centrosomal protein of 85 kDa | s34959 / s34961 | 5 | correlated / anti-correlated | 0.64 |
| **CTSB** | Cathepsin B | s3738 / s3739 | 1 | anti-correlated | 0.53 |
| **DSE** | Dermatan Sulfate Epimerase | s26749 / s26750 | 1 | anti-correlated | 0.54 |
| **EME1** | Essential Meiotic Structure-Specific Endonuclease 1 | s44946 / s44945 | 1 | correlated | 0.66 |
| **ESYT2** | Extended synaptotagmin-2 | s33138 / s33136 | 2 | anti-correlated | 0.75 |
| **F11R** | F11 Receptor | s27152 / s27151 | 6 | anti-correlated | 0.55 |
| **FOXM1** | Forkhead Box M1 | s5250 / s5249 | 1 | correlated | 0.68 |
| **LGR4** | Leucine-Rich Repeat G Protein-Coupled Receptor 4 | s30840 / s229314 | 1 | anti-correlated | 0.61 |
| **LHFP** | Lipoma HMGIC Fusion Partner | s19847 / s19848 | 1 | correlated | 0.57 |
| **LMNB2** | Lamin B2 | s39477 / s39476 | 1 | anti-correlated | 0.60 |
| **LRP1** | LDL Receptor Related Protein 1 | s8278 / s8280 | 4 | anti-correlated | 0.70 |
| **MAP7** | Ensconsin | s17263 / s17262 | 2 | correlated | 0.67 |
| **MEIS2** | Meis Homeobox 2 | s8666 / s8664 | 7 | correlated / anti-correlated | 0.57 |

| | | | | | |
|---|---|---|---|---|---|
| **MYC** | Myc proto-oncogene protein | s9130 / s9131 | 2 | anti-correlated | 0.68 |
| **PLK1** | Polo-like kinase 1 | s448 / s450 | 4 | correlated | 0.64 |
| **PRC1** | Protein regulator of cytokinesis 1 | s17268 / s17269 | 1 | anti-correlated | 0.73 |
| **RAN** | GTP-binding nuclear protein Ran | s11769 / s11768 | 1 | anti-correlated | 0.60 |
| **RBBP4** | Histone-binding protein RBBP4 | s55169 / s56872 | 1 | anti-correlated | 0.57 |
| **RGMA** | Repulsive Guidance Molecule Family Member A | s32498 / s32500 | 7 | correlated / anti-correlated | 0.70 |
| **TCF7** | Transcription factor 7 | s13877 / s13878 | 2 | anti-correlated | 0.71 |
| **TOP2A** | Topoisomerase II Alpha | s14307 / s14308 | 2 | correlated | 0.66 |
| **TUFT1** | Tuftelin | s14510 / s14509 | 2 | anti-correlated | 0.59 |
| **WBP1** | WW Domain Binding Protein 1 | s24095 / s225969 | 1 | correlated | 0.52 |

**Supplementary Table 2: Comparison of spinning disc and diSPIM microscopy**

Comparison of the acquisition properties and resulting image quality between a spinning disc microscope (Zeiss LSM 780) and the diSPIM system. Light-sheet imaging outperforms spinning disc microscopy in resolution, acquisition speed, signal-to-noise ratio and phototoxicity.

| | Spinning disc microscope | diSPIM |
|---|---|---|
| **XYZ stack (px x px x slices)** | 1004 x 1002 x 233 | 2x (1024 x 1024 x 260) |
| **resolution** | 0.2 µm / px | 0.1625 µm / px |
| **laser power** | 1,320 µW / s | 320 µW / s |
| **exposure / slice** | 50 ms | 1.75 ms |
| **stack acquisition duration** | 53.2 s | 4.5 s (+ 15 s) |
| **signal-to-noise ratio** | 43.25 | 127 |
| **avg. background signal** | 18.4 | 0.941 |
| **power density / phototoxicity** | 168,000 $W/cm^2$ | 40,700 $W/cm^2$ |

**Supplementary Table 3: 23 features describing spheroid phenotypes**

| Feature | Name | Description | Global or nuclear feature |
|---|---|---|---|
| **1** | **spheroid growth rate (nuclei)** | rate of increase in nuclei count over the course of the time lapse | global |
| **2** | **prophase ratio** | fraction of nuclei classified as "prophase" | nuclear |
| **3** | **metaphase ratio** | fraction of nuclei classified as "metaphase" | nuclear |
| **4** | **anaphase ratio** | fraction of nuclei classified as "anaphase" | nuclear |
| **5** | **avg. cell volume** | average cell volume (in voxels) as ratio of spheroid volume to nuclei number | global |
| **6** | **prophase segment volume** | average nucleus size (in voxels) across all nuclei classified as "prophase" | nuclear |
| **7** | **metaphase segment volume** | average nucleus size (in voxels) across all nuclei classified as "metaphase" | nuclear |
| **8** | **anaphase segment volume** | average nucleus size (in voxels) across all nuclei classified as "anaphase" | nuclear |
| **9** | **interphase segment volume** | average nucleus size (in voxels) across all nuclei classified as "interphase" | nuclear |
| **10** | **spheroid volume** | spheroid volume (in voxels) throughout the time lapse | global |
| **11** | **avg. segment volume** | average nucleus size (in voxels) across all nuclei in all cell cycle phases | global |
| **12** | **spheroid growth rate (volume)** | rate of volume increase of the spheroid hull throughout the time lapse | global |
| **13** | **spheroid compactness** | factor describing the volume in relation to the largest extent | global |
| **14** | **convexity** | factor describing the volume in relation to the surface area | global |
| **15** | **nuclei migration speed** | average movement of all nuclei in 3D space in pixel per time point | global |
| **16** | **interphase transition duration** | average duration a nucleus spends in "interphase" | nuclear |
| **17** | **prophase transition duration** | average duration a nucleus spends in "prophase" | nuclear |
| **18** | **metaphase transition duration** | average duration a nucleus spends in "metaphase" | nuclear |
| **19** | **anaphase transition duration** | average duration a nucleus spends in "anaphase" | nuclear |
| **20** | **total number cell cycle transitions** | total number of deduced cell cycle phase transitions | global |
| **21** | **normal / abnormal transition** | fraction of cell cycle phase transitions that are biologically implausible | global |
| **22** | **spheroid roundness** | factor describing shape of spheroid | global |
| **23** | **size / spheroid roundness ratio** | ratio of spheroid volume to roundness | global |

### Methods and Materials

#### Methods

##### Culture of MCF10A H2B-GFP cells

MCF10A H2B-GFP cells (passage 25 to 31) were cultured in 2D in 25 $cm^2$ culture flasks (Greiner bio-one) in DMEM/F12 medium (ThermoFisher Scientific #11039) with supplements (5% horse serum, 10 µg/ml Insulin (Life Technologies), 20 ng/ml EGF, 0.5 mg/ml hydrocortisone and 100 ng/ml Cholera Toxin (Sigma)) under standard culture conditions (5% $CO_2$ / 37 °C), and passaged after reaching 80-90% confluency with 0.05% Trypsin (Life Technologies) every three to four days.

##### Solid-phase reverse siRNA transfection

Solid-phase reverse transfection siRNA transfection mix was prepared as described (*22*), but using trehalose dihydrate (Merck #T9531) instead of sucrose.
For transfection, trypsinated MCF10A H2B-GFP were diluted in growth medium to a density of $5x10^5$ cells/ml. 10'000 cells in 100 µl cell suspension were added to each well of the solid-phase reverse transfection mix. After five hours, cell medium was removed and cells were resuspended by directly adding 50 µl 0.25% Trypsin (Life Technologies #25200056) to each well.

##### High-content cell spotting in Matrigel

Mixing of cells with Matrigel (Corning Matrigel Matrix) and spotting into OneWell plates (Greiner bio-one CELLSTAR® OneWell Plate™ #670180) was conducted by an automated liquid handling robot from Hamilton Robotics with a custom protocol. In short, from each cell suspension transfected with individual siRNA, 60 isolated cells in 3 µl medium were mixed with 10 µl Matrigel. Subsequently each mixture was spotted eight times with a single spot volume of 0.2 µl in a two columns by four rows array, resulting in a total of 320 spots in 40 columns and eight rows on the imaging plate. One sub-array of spots was always dedicated to beads (ThermoFisher Scientific #7220) mixed with Matrigel, used for registration of the two acquired views. Positioning of each spot is identical with the positions of a standard 1536-well plate. After 10 minutes at 37°C for Matrigel solidification, culture medium was added and samples were incubated under standard culture conditions until imaging.

##### diSPIM imaging

Imaging was conducted with a dual-view inverted selective plane illumination microscope (diSPIM) as described (*4*). The microscope was equipped with LMM5 laser (Spectral Applied Research Laser Illumination Laser Merge Module 5) and AHF Quad Filterset (F59-405 / F73-410 / F57-406). Images were acquired by two water-cooled ORCA-Flash4.0 Hamamatsu sCMOS cameras. Cooling was provided with Julabo F250 cooling circuit. Standard culture conditions were provided by an incubation chamber (3i ECS2) and direct airflow over the SPIM head was minimized to avoid unnecessary vibrations. All imaging time lapse acquisitions were conducted with 320 µW laser power for 488 nm excitation wavelength (measured at the sample). Readjusting the fine alignment of the microscope was conducted shortly before the start of the acquisition.

**Low resolution pre-screen**

To detect each spheroid's positions and select the spheroids to be imaged, we conducted a fast, low resolution stage-scan pre-screen. A grid of imaging positions was defined across the imaging plate, with each position placed at the center of one column of spots. As the automated spotting process resulted in spots with defined positions and sizes, we were able to repeatedly use the same grid of stage scan acquisition positions for every pre-screen. Potentially due to small manufacturing differences of the imagining plate, we solely needed to adjust the general Z-position off-set, underlining the robustness of our sample preparation process. Each position acquisition resulted in a $X_{microscope}$-Stack of 1 200 slices with a step size of 5 µm, a pixel resolution of 0.648 µm/px and a field of view of 333 µm.

The acquisition of the pre-screen took 31 minutes and produced 96 000 images. This pre-screen data was subsequently analysed by a KNIME image processing workflow detecting the $XYZ_{microscope}$ position, size and shape of each cell cluster. Per spot, we detected an average of 2.6 spheroids. Small, flat and elongated cell clusters were excluded and the remaining spheroids ranked based on their Z-position. To minimize obstructions in the illumination and detection path and maximize image quality, the spheroids with the largest Z-coordinates were selected for imaging for each condition.

38 defined cell spheroids plus two positions with fluorescent beads (used for image processing) were imaged with the diSPIM microscope for 24 hours at maximal temporal and spatial resolution for treatment evaluation.

**Position scan acquisition**

Preselected positions from the KNIME analysis of the pre-screen were checked and if necessary manually corrected. An additional registration position was added as first and last position, imaging beads mixed in Matrigel.

Imaging parameters for dual view synchronous piezo/slice scan (stack acquisition) were set to acquire two stacks of 1024 $px^2$ in $XY_{image}$ with the maximal camera resolution of 0.1625 µm/px centered to the field of view of the camera and 260 slices in $Z_{image}$ with a slice interval of 0.5 µm starting with view A (right camera acquisition). Sample exposure was set to 1.5 ms and the option for "Minimized Slice Period" enabled. The option for "Autofocus during acquisition" was enabled with the autofocus running on the registration position imaging beads every acquisition cycle with 40 slices acquired every 0.5 µm. The off-set was detected by the "Vollath" algorithm.

Due to the acquisition limitations of the microscope of a minimal four to five seconds per position scan and stage repositioning, we imaged at an interval of five minutes for 24 hours, resulting in a total data volume of 10.06 terabytes (TB), which was stored locally. The high temporal resolution was essential for tracking of nuclei as they progressed through the cell cycle. Throughout time lapse acquisition, we did not need to adjust for any position off-set introduced by deformation of the matrigel or external influences as samples remained almost universally in the field of view.

**Image processing (hSPIM)**

Raw data was processed by a custom software named 'hSPIM' specifically adapted to the geometry of the diSPIM and the separately acquired registration beads positions. In hSPIM, the registration matrix of the two views is detected for each time point of the screen by registration of beads in 3D. Additionally, the PSF is extracted. This registration matrix and PSF are stored and used for registration and deconvolution of all other acquired positions at this time point. Furthermore, the software performs a segmentation of the nuclei, from which different geometrical and textural features are extracted for each segment. Deconvolved fusion images and segment images as well as segment and feature table are stored and were used for further image analysis. In addition, the hSPIM software can directly visualize in 3D a registered and deconvolved image snapshot, store a view angle, and export a 3D movie of a single position. Library code and documentation for hSPIM are available at *https://github.com/eilslabs/diSPIM_screen*.

**High-content KNIME analysis workflow**

Following the raw image processing, we developed a KNIME workflow to analyze key cellular and global properties of each spheroid throughout the acquired time lapse. The workflow is available at *https://github.com/eilslabs/diSPIM_screen*.

***$XYZ_{microscope}$ displacement****:* By tracking the positions of single beads over time, we could detect and correct for the global offset in all dimensions of the microscope introduced through fine displacements of the imaging plate or expansion of the diSPIM components.

**Clustering of segments into spheroids**: To segregate segments from two spheroids acquired at a single imaging position into individual spheroid clusters, we analyzed the geometric distance of each segment to all others and clustered segments accordingly.

**Spheroid size:** The clustering enabled us to combine all segments of one spheroid and determine spheroid size.

***Cell cycle phase classification****:* For precise cell cycle phase classification, we used a VGG-based convolutional neuronal network trained on a set of manually classified images. The CNN calculated the probability for each of the four cell cycle phases (interphase, prophase, metaphase, anaphase) for each XY, XZ and YZ slice. The class with the highest sum in likelihood for each segment was selected as the cell cycle phase of the nucleus at this time point.

***Geometric nuclei class features****:* Single nuclei size, intensity, position of segments from the center of the spheroid and predicted cell cycle class were recorded over time.

***Nuclei migration speed****:* By tracking the position of each segment over time, we analyzed the median migration speed of all cells in each spheroid.

***Time lapse movie****:* For individual evaluation, we exported the maximum projected time lapse movie of each position, including cell cycle classification and spheroid hull.

**Spheroid feature evaluation**

Selected image features quantified by the KNIME workflow (Supplementary Table 3) were subjected to further quantitative analysis in R. Additional quantitative features such as mean cell volume (estimated as the ratio of spheroid volume to nuclei number) and average nuclei size in different cell cycle phases were computed. The fraction of cells detected in different cell cycle phases was averaged across all time points. To calculate instantaneous spheroid growth rates from nuclei numbers, the number of nuclei over time was smoothed using the lowess function with parameters f=1/3, iter=3L, delta=0.01 * diff(range(NrCells[1:n.rows[[pos]],pos])), and differentiated using the diff function.

All feature measurements from all plates were combined into one matrix, centered by subtracting the column means from their corresponding columns, and scaled by dividing the centered columns by their standard deviations. As mechanical plate drift resulted in spheroids lying partially outside the field of view in one plate, affected features (nuclei and spheroid growth rate, compactness, convexity, sphericity, spheroid volume and cell volume) were excluded for this plate.

To identify clusters of siRNAs causing similar phenotypes, rank-based clustering was performed using the rank, dist, and hclust functions. Heatmaps were created using the heatmap.2 function from the gplots package or the aheatmap function from the NMF package.

**dCas9-effector domains construct synthesis**

We fused the catalytic C-terminal effector domains of epigenome modifying enzyme (DNMT3a, TET1) C-terminally to the dCas9 via a linker and added two nuclear localization sequences (NLS) for improved nuclear localization and a M2 flag to the N-terminus. A dCas9 with no C-terminal addition of an ED was used as binding control and to physically block binding sites for regulatory factors. Source constructs were obtained from AddGene for dCas9 (*31*), DNMT3a (*33*) and TET1 (*34*). dCas-ED constructs (C49 – dCas9; C54 – DNMTA3-dCas9; C57 – TET1-dCas9) were assembled by Gibson Cloning (NEB #E2611) following manufacturer guidelines. Linker (GGGGS), NLS (PKKKRKV) and M2-Flag (DYKDHDG) DNA sequences as well as adapter primers were ordered from Eurofins Genomics. Successful cloning was assessed by sequencing by GATC Biotech AG, western blot (M2-flag) and expression in HEK293 cells detected by immunostaining. Plasmid maps and construct components are available from the authors on request.

**Stable dCas9-ED expression in HEK293 cell line**

$5 \times 10^5$ HEK293 cells were transfected with 5 µg plasmid DNA of the different dCas9-ED constructs (C49, C54, C57) with Lipofectamin 2000 (Invitrogen) following manufacturer guidelines. 48 hours post dCas9-ED plasmid transfection, transfected cells were selected by addition of G418 (Geneticin) antibiotic to the culture medium at a concentration of 500 µg / ml. Stable dCas9-ED expressing cell lines were frozen after four passages under G418 selection.

**CpG selection**

We targeted the different epigenome modifying molecular tools to specific genomic sites by combining different sgRNAs with the different effector domains to modify distal or proximal regulatory CpGs with regulatory properties (Supplementary Table 1). Based on a data set comprising 450k Illumina gene expression and CpG methylation data from human breast cancer patients (*35*) available on the UCSC genome browser (*36*), we selected CpGs with high correlation (Pearson correlation > 0.5) or high anti-correlation correlation (Pearson correlation < -0.5) between CpG methylation level and target gene expression. We detected up to seven correlated or anti-correlated regulatory CpGs per target gene. Correlated CpGs (low CpG-me results in reduced expression) are expected to result in a gene knock-down phenotype when targeted by the TET1 dCas9-ED, while anti-correlated CpGs (high CpG-me results in reduced expression) are expected to show the abnormal mitotic phenotype when targeted by DNMT3A. Target genes that had only a single regulatory CpG with a correlation between expression level and CpG methylation below 0.6 were not further analyzed, which excluded ATHOH8, AURKA, BUD31, CTSB, DSE, ESYT2, LGR4, RAN, and RBBP4.

**Single guide RNA design and synthesis**

sgRNAs directing the dCas9 effector domain fusion protein to the specific genomic site were designed to direct the methylome-modifying enzymes to positions around 33 base pairs upstream from their corresponding target CpG, since the dCas9-ED has been described to show highest epigenome modifying effectivity at 27 bp (+/-17 bp) from the PAM sequence of the sgRNA (*33*). We designed two opposing sgRNAs per regulatory CpG, one binding to the sense and one binding to the anti-sense strand of the DNA. Furthermore, sgRNA target sites had a minimum of two mismatches to the next off-target site, to reduce off-targeting effects.

To evaluate gene knock-down through binding of the dCas9 without added effector domain to the transcription start site (TSS), we used the FANTOM5/CAGE online atlas (http://fantom.gsc.riken.jp/5/) to define the TSS of our target genes and selected a single sgRNA binding site at an average of 50 bp upstream of the TSS for optimal gene repression (*37*, *38*).

sgRNA expression plasmids were designed and synthesized following a previously published SAM target sgRNA cloning protocol (*39*). In short, the sgRNA(MS2) cloning backbone (AddGene #61424) was digested with BbsI. Oligos representing the sgRNA target site with 20 bases in sense (Os) with a CACCG overhang and anti-sense (Oas) with an AAAC overhang were ordered from Eurofins and annealed. For genome reference, we used the UCSC Genome Browser on Human Feb. 2009 (GRCh37/hg19). Backbone and sgRNA defining insert were joined by a Golden Gate reaction. The resulting plasmid was expanded by bacterial transformation and assessed by sequencing.

**Stable dCas9-ED cell lines sgRNA transfection**

The HEK293 cells were transfected with the different sgRNA constructs by solid-phase reverse transfection as described (*22*) but using Lipofectamine 2000 (Invitrogen #11668027) instead of Lipofectamine RNAiMAX.

**Immunostaining of HEK293 cells for DNA, sgRNA and dCas9-ED**

HEK293 cells were fixed and stained at different time points between 3 and 9 days after transfection for the two components of the functional dCas9-ED by immunofluorescence (IF) staining. Cells were fixed with 4% PFA (Sigma-Aldrich #F8775) for 10 minutes in PBS with 0.5% Triton X-100 and blocked subsequently with 1% goat serum in PBS applied overnight. Mouse anti-Flag M2 monoclonal primary antibody (Sigma #F1804) and goat anti-mouse Alexa 568 secondary antibody (Invitrogen #A11004) were used to label the dCas9-ED. Successful transfection with the sgRNA plasmid was detected with rabbit anti-GFP monoclonal primary antibody (Cell Signaling #2956) and goat anti-rabbit Alexa 488 (Molecular Probes #A11034). DNA was stained with DAPI.

**Confocal imaging of IF stained epigenome targeted HEK293 cells**

Confocal imaging was conducted using the Zeiss LSM 780 with the AutofocusScreen macro (http://www.ellenberg.embl.de/apps/AFS/, 24.02.2016), acquiring 25 Z-stacks per well with each comprising five slices per dCas9-ED-sgRNA combination (one dCas9-ED / one target gene). Each stack was acquired with a bright field image additionally to the DAPI (405 nm), sgRNA (488 nm) and dCas9-ED (568 nm) channels.

**2D pre-screen of dCas9-ED HEK293 cells and phenotype evaluation**

In a 2D pre-screen designed to select for significant methylome regulated target genes, a total of 129 possible dCas9-ED-sgRNA combinations were evaluated with an average of 6856 cells analyzed per combination. Solid-phase reverse transfection was used to deliver the sgRNA expressing plasmid into the dCas9-ED expressing cell lines, where its expression was confirmed by GFP expression. We evaluated and classified the nuclear phenotype of cells expressing dCas9-ED and the sgRNA at 3, 5 and 7 days post transfection.

Raw HEK293 images of each sgRNA-dCas9-ED combinations were smoothened by Gaussian convolution and single nuclei were segmented by Otsu thresholding. Single segments were further processed and split if necessary by segment erosion. Using the same CNN architecture as above, this time trained on annotated images of 2D HEK293 cells, single nuclei were classified by into cell cycle stages (inter-, pro-, meta-, anaphase) as well as significant phenotypes (macronuclei and apoptotic condensed DNA). Additionally, the transfection state of the cell was evaluated by the presence of dCas9-ED (M2-flag) and sgRNA (GFP), and only cells expressing both components were included in the analysis.

Detected classes were further evaluated in comparison to non-targeted sgRNA transfected dCas9-ED cell lines as well as to non-transfected cells. All acquired time points were combined during analysis. Cells with a significantly higher (> 1.5 fold) occurrence of a class compared to control cells were highlighted.

We found that only 18% of sgRNA-dCas9 combinations showed a significant effect on the mitotic phenotype, although 85% of those phenotypes correlated with the siRNA induced phenotype and 88% correlated with knock-down phenotypes published in the online databases MitoCheck (*23*) and Cyclebase (*40*) (Supplementary Figure 5). The targeted CpG properties matched with the expected gene regulatory effect of the dCas9-ED in only 8/18 cases, suggesting that the majority of abnormal mitotic phenotypes were evoked by the dCas9 protein blocking access to regulatory sites.

#### Materials

##### Hardware

*Workstation*

| hardware | supplier | description |
|---|---|---|
| CPU | Intel | i9-7980XE |
| GPU | NVIDIA | Titan xp 12 GB |
| Hard drive (RAID0) | WD | WD-Red 8 TB |
| RAM | ECC | 64 GByte DDR-4 PC2400 |
| Motherboard | ASRock | X299 Taichi |
| Controller | Intel | SATA Controller, 10x 6 Gbit/s |
| Hard drive | Samsung | 1 TB 960 Pro |

*ASI diSPIM hardware*

| hardware | supplier | description / number |
|---|---|---|
| camera cooling | Julabo | F250 |
| quand filterset | AHF | F59-405<br>F73-410<br>F57-406 |
| sCMOS cameras | Hamamatsu | ORCA-Flash4.0 |
| laser | Spectral Applied Research | Laser Merge Module 5 (LMM5) |

##### Software and workflows

*Software*

| name | version | description |
|---|---|---|
| KNIME | 3.5.5 | Konstanz Information Miner |
| hSPIM | 1.0 | diSPIM raw image processing tool ('hSPIM')* |
| MicroManager | 1.4 | microscope control software |
| diSPIM plugin | NB_20180116 | nightly build MicroManager diSPIM controller plugin |

*hSPIM library available at *https://github.com/eilslabs/diSPIM_screen*

*KNIME workflows (available at https://github.com/eilslabs/diSPIM_screen)*

| diSPIM_prescreen_stagescan_Pos_analysis |
|---|
| diSPIM_phenotype_screen_analysis_3D_spheroids |
| EpiTool_confocal_nuclei_classification |
| EpiTool_Class_quantitative_analysis |

*Haralick features used for phenotype characterization*

| Haralick feature | description | Haralick feature | description |
|---|---|---|---|
| F1 | contrast | F8 | sum entropy |
| F2 | angular second moment | F9 | entropy |

| | | | |
|---|---|---|---|
| F3 | correlation | F10 | difference variance |
| F4 | sum of squares: variance | F11 | difference entropy |
| F5 | inverse difference moment | F12 | measure of correlation I |
| F6 | sum average | F13 | measure of correlation II |
| F7 | sum variance | | |

**Source constructs**

| construct | source | description / number |
|---|---|---|
| #46911 | AddGene | Gilbert_pHR-SFFV-dCas (*31*) |
| #71666 | AddGene | pdCas9-DNMT3A-EGFP (*33*) |
| #49792 | AddGene | FH-TET1-pEF (*34*) |
| #61424 | AddGene | sgRNA(MS2) cloning backbone |

**Antibodies**

| description | source | number | description |
|---|---|---|---|
| anti-Flag® M2 | Sigma | F1804 | primary mouse anti Flag M2 monoclonal antibody |
| anti-GFP | Cell signaling | 2956 | primary rabbit anti GFP monoclonal antibody |
| Anti-rabbit Alexa 488 | Molecular Probes | A11034 | fluorescent secondary goat anti rabbit antibody |
| Anti-mouse Alexa 568 | Invitrogen | A11004 | fluorescent secondary goat anti mouse antibody |

**Consumables and solutions**

| description | supplier | product number |
|---|---|---|
| beads: PS-Speck™ Microscope | ThermoFisher Scientific | P7220 |
| Cell Culture Plate, 96-Well | Eppendorf | 0030730119 |
| CELLSTAR® OneWell Plate™ | Greiner bio-one | 670180 |
| Cholera toxin | Sigma-Aldrich (Merck) | |
| Collagen type IV solution | Merck | C5533 |
| culture flasks (25cm$^2$) | greiner bio-one | |
| DAPI | Sigma-Aldrich (Merck) | D9542 |
| DMEM/F12 | ThermoFisher Scientific | 11039 |
| G418 (Geneticin) | Sigma-Aldrich (Merck) | 4727878001 |
| Gibson Assembly Master Mix | NEB | E2611 |
| Insulin | Life Technologies | |
| Lipofectamine 2000 | Invitrogen | 11668027 |
| Lipofectamine® RNAiMAX | ThermoFisher Scientific | 13778075 |
| Matrigel | Corning | 354248 |
| OptiMEM | ThermoFisher Scientific | 51985026 |
| PCR plate, 96 well | Kisker | G060 |
| trehalose dihydrate | Merck | T9531 |
| trypsin | Life Technologies | 25200056 |
